# Acclimation temperature and parasite infection drive metabolic changes in a freshwater fish at different biological scales

**DOI:** 10.1101/2024.06.28.599683

**Authors:** Marie Levet, Shaun S. Killen, Stefano Bettinazzi, Vincent Mélançon, Sophie Breton, Sandra A. Binning

**Affiliations:** Département de Sciences Biologiques, Université de Montréal, 1375 Avenue Thérèse-Lavoie-Roux, Montréal, Québec, Canada, H2V 0B3; Groupe de recherche interuniversitaire en limnologie (GRIL), Université de Montréal,1375 Avenue Thérèse-Lavoie-Roux, Montréal, Québec, Canada, H2V 0B3; School of Biodiversity, One Health and Veterinary Medicine, Graham Kerr Building, University of Glasgow, Glasgow, Scotland, United Kingdom, G12 8QQ; Department of Genetics, Environment, and Evolution, University College London, 99-105 Gower Street, London WC1E 6BT, United Kingdom

## Abstract

1. Environmental stressors such as elevated temperature and parasite infection can impact individual energy metabolism. However, organismal responses to co-occurring stressors and their effects across biological scales remain unexplored despite the importance of integrative studies for accurately predicting the resilience of natural populations in changing environments.
2. Using wild-caught, naturally parasitized pumpkinseed sunfish, Lepomis gibbosus, we quantified changes in cellular and whole-organism metabolism in response to temperature and parasite infection. We acclimated pumpkinseeds for three weeks at 20°C, 25°C, or 30°C before measuring whole-organism oxygen uptake (ṀO2) using intermittent flow-respirometry to quantify maximal and standard metabolic rates (MMR and SMR, respectively) and aerobic scope (AS). We also measured the maximal activity of enzymes (citrate synthase (CS), respiratory complexes I + III and IV of the electron transport system, and lactate dehydrogenase (LDH)) linked with cellular bioenergetics in fish heart, brain, spleen and gills using spectrophotometry.
3. We found no interactions between acclimation temperatures and parasite intensity on cellular or whole-organism metabolism. However, both stressors were independently related to fish metabolism, with differing impacts across biological scales.
4. Whereas MMR increased with acclimation temperature, this was not mirrored by increasing SMR or decreasing AS, suggesting thermal compensation across acclimation temperatures at the whole-organism level.
5. On a cellular level, acclimation responses were similar across organs, with maximal activity of all enzymes decreasing with increasing acclimation temperature. However, LDH activity remained higher than aerobic enzyme activities (CS, ETS complexes I + III and IV) across acclimation temperatures and organs, especially in gills, where LDH activity drastically increased at 30°C. This may indicate a stronger reliance on anaerobic metabolism to sustain whole-organism metabolic performance.
6. Fish with greater trematode infection had lower MMR and AS. There were no relationships between parasite intensity and SMR nor maximal enzyme activity.
7. Our work shows that co-occurring stressors have distinct impacts on fish metabolism: parasites are primarily related to whole-organism metabolism while temperature impacts metabolism across biological scales. This highlights that interactions among co-occurring stressors are important for ecological realism and accurate predictions regarding population resilience to environmental changes.

## Introduction

Warming water temperatures are threatening freshwater ecosystems globally (Hassan et al., 2020). Ectotherms, such as freshwater fishes, are sensitive to these shifts in ambient temperature. Warmer waters can alter the rate of cellular and whole-organism level processes, ultimately affecting an individual’s capacity to perform in their environment (Guderley & St-Pierre, 2002; Schulte, 2015). To counteract the direct effects of temperature, fishes can thermally acclimate through physiological adjustments (e.g. changes in enzyme activities, organ morphology, oxygen uptake) to enable sufficient ATP production for the maintenance of metabolic performance across thermal environments (Schulte, 2015). Although the physiological responses underlying thermal acclimation are well-studied in some species and habitats, gaps in our understanding remain (Jutfelt et al., 2024).

Most studies investigating metabolism during acclimation concentrate either on changes at the organismal or sub-organismal levels. However, evidence indicates that metabolic responses might vary across levels of biological organisation (Iverson et al., 2020). There is thus an urgent need for research exploring metabolic rate measurements at different scales to better understand how ectotherms cope with variations in their environment. At the whole-organism level, elevated temperatures can increase an organism’s standard metabolic rate (SMR, i.e. the minimum energy required to maintain basal physiological demands at a given temperature) and maximum metabolic rate (MMR, i.e. the maximum rate of aerobic energy expenditure). Variations in SMR and MMR directly affect an organism’s aerobic scope (AS), calculated as the difference between SMR and MMR, which represents an organism’s capacity to aerobically perform fitness-enhancing activities that require energy, such as growth, digestion, and reproduction (Fry, 1971).

At the sub-cellular level, acclimation to temperature involves quantitative changes in the production and activity of enzymes (Somero, 2004). In response to elevated temperature, the activity levels of aerobic and anaerobic metabolic enzymes, such as citrate synthase (CS) and lactate dehydrogenase (LDH), may be altered to meet the increased energy demands associated with metabolic adjustments triggered by temperature changes (Grim et al., 2010; Ressel et al., 2022). Enzymes related to the respiratory chain are also subject to similar changes. For instance, studies on warm-acclimated fishes found that rainbow trout (*Oncorhynchus mykiss*) decrease CCO activity as acclimation temperature increases and common killifish *(Fundulus heteroclitus*) acclimated to high temperatures (33°C) have altered ETS performance (Chung et al., 2017; Kraffe et al., 2007). Variation in reaction rates can also occur among organs in response to thermal acclimation (Chung, Bryant, and Schulte 2017; Cominassi et al. 2022). For example, in the skeletal muscle of threespine stickleback (*Gasterosteus aculeatus*), the maximal activity of CS and LDH was equivalent among acclimation groups (5°C, 12°C, and 20°C) (Cominassi et al., 2022). However, stickleback acclimated at 20°C showed higher maximal activity of CS and LDH in the liver compared to fish acclimated at 5°C and 12°C. Despite mounting evidence suggesting organ-specific responses to acclimation, studies looking for general patterns across many organs remain scarce. Such studies are especially important for work on wild animals, where organs may differ in their response to acclimation because of interacting effects with biotic stressors.

Among biotic stressors, parasites are a critical part of all ecosystems but are often an overlooked ecological component despite their pervasive effects on host performance (Chrétien et al., 2023; McElroy & Buron, 2014). Parasite infections can alter host metabolic rates, organ mass, and the availability of metabolic substrates (Nadler et al., 2021; Ryberg et al., 2020). Furthermore, recent studies on wild-caught parasitized pumpkinseed sunfish (*Lepomis gibbosus*) have revealed substantial variability in metabolic enzyme activity across different organs (Sabbagh et al. *In Press*; Mélançon et al. 2023). Parasites may also influence individual acclimatory responses to temperature. For instance, the bacterial pathogen *Pasteuria ramosa* reduced the heat tolerance of the *Daphnia magna* host (Hector et al., 2024). To date, no studies on wild animals have included parasites in investigating the physiological responses to thermal acclimation across different biological scales. Therefore, whether and how parasitism relates to an organism’s response to temperature acclimation remains an open question.

Here, we explored how thermal acclimation and parasitism affect the metabolism of pumpkinseed sunfish (*Lepomis gibbosus*; Linnaeus, 1758) at multiple levels of biological organisation (cellular and whole-organism). Pumpkinseeds host several endoparasite species, including cestodes, nematodes and trematodes (e.g. *Uvulifer sp., Apophallus sp.,* i.e. black-spot disease) (Margolis & Arthur, 1979). Our study addressed the following questions: (1) Does thermal acclimation and/or parasites alter whole-organism metabolic rates? (2) Does metabolic enzyme activity vary in response to thermal acclimation and/or parasites? (3) If so, are there any organ-specific patterns in enzymatic activity related to temperature and/or infection? At the whole-organism level, we expected increased whole-organism metabolic rates in response to thermal acclimation. At the same time, we predicted a negative relationship between whole-organism metabolic rates and parasite intensity. At the cellular level, we expected decreased enzymatic activity with increasing acclimation temperature and parasite intensity. If both acclimation temperature and parasite intensity influence host metabolism, then we expected an interactive effect of increased temperature and parasite intensity.

## Materials and Methods

### Fish collection and husbandry

In June 2022, we captured 120 wild pumpkinseed sunfish (mean ± s.d. Total length = 8.95 ± 0.74 cm, body mass = 12.64 ± 3.16 grams) in Lake Cromwell located near the Université de Montréal’s Station de biologie des Laurentides (Canada, 45.98898°N, −74.00013°W). Fish were captured with baited minnow traps placed in the lake’s littoral zone. Selected fish were transferred in opaque containers filled with lake water aerated by a transportable air bubbler. Upon arrival at the SBL laboratory housing facilities (15 min walk), fish were placed in a saltwater bath (3 g/L) to reduce the risk of fungal and bacterial infections. All salt bath tanks were 40 litres (10 -15 fish per tank), with water maintained at 18°C (i.e. water temperature at capture). Water was aerated, changed and siphoned twice a day to maintain water quality. Post-quarantine, fish were weighed, measured and uniquely tagged with visible elastomer implants (VIE; Northwest Marine Technology) on both sides of the dorsal fin with a 29-gauge needle. Fish were distributed among three 600L flow-through holding tanks (215 × 60 × 60 cm, length × width × height) with 40 fish per tank, separated into three sections in which artificial plants and PVC refuges were used for environmental enrichment. Water entering the flow-through tanks was sourced from a nearby lake (Lake Croche; 45.99003°N, −74.00567°W) with a water profile similar to Lake Cromwell. Water was particle-filtered and UV-sterilised before entering the tanks. The light photoperiod followed the summer light-dark cycle (14L:10D). Fish were fed ad libitum daily with bloodworms.

### Acclimation treatments

Environmentally relevant acclimation temperatures were selected based on water temperature data collected from June to September 2021 with a temperature logger (HOBO Pendant MX 2201, USA) placed at a 1-meter depth in the littoral zone of Lake Cromwell (Fig. S1): 20°C (early-summer temperature), 25°C (mid-summer temperature), and 30°C (climate warming projection), This set this temperature 2°C higher than the highest temperature recorded during the summer of 2021 (Fig. S1). We increased water temperature in holding tanks 24 hours after the tagging procedure (1.5°C day ^−1^). Temperature was maintained with heating (1500W and 2000W heaters, GESAIL) and cooling systems (EK20 immersion cooler, Thermo ScientificTM, USA) connected to a temperature controller (ITC-308S, Inkbird) (Fig. S2). We also monitored the water temperature twice a day with a digital thermometer (Hanna HI98509 *Checktemp 1* Digital Thermometer). Each tank had a constant inflow of oxygenated water with a replacement rate ranging from 4.8L/h to 22.8L/h. Air bubblers throughout the tank ensured air saturation, with submersible pumps (ECO-396, EcoPlus®, China) for water mixing. Fish were acclimated for at least three weeks to allow thermal compensation (Dent & Lutterschidt, 2003).

### Whole-organism metabolic rates

We measured the fish’s oxygen uptake rate (*Ṁ*O2; mg O2 ⋅h^-1^) using intermittent-flow respirometry. We followed the guidelines described by Killen et al. (2021). Each day, we measured between 8 and 12 fish in individual acrylic respirometry chambers placed in 80L water-controlled baths. Detailed information regarding water-recirculation system and temperature control can be found in Table S1. Each respirometry chamber was opaque, with a top viewing window to minimise disturbances. A Firesting fibre optic oxygen meter (Firesting 4-channel oxygen meter, PyroScience GmbH, Aschem, Germany) located in the recirculation loop measured the dissolved oxygen concentration every 3 seconds.

Fish were fasted for 24 hours before the start of the trial. Each trial started between 15:00 and 18:00, with an estimate of the maximum metabolic rate (Peak *Ṁ*O2recovery; hereafter MMR). Each fish was placed in a circular arena (31 cm diameter) filled with 8L of water from the holding tank. Fish were manually chased for three minutes and exposed to air for one minute before being transferred to the respirometry chamber (Roche et al., 2013). *Ṁ*O2 was continuously measured for ∼ 15 minutes to estimate MMR, defined as the upper limit of the ability of the fish to perform aerobic metabolism. Afterwards, fish remained in the chamber between 19-22h to estimate their standard metabolic rate (*Ṁ*O2min; hereafter SMR), defined as the minimum energy required to maintain basal physiological demands.

MMR was analysed using the respR package in R (Harianto et al., 2019). We used the rolling regression method with a two-minute window width over ∼ 15 minutes to determine the highest rate of *Ṁ*O2 (Prinzing et al., 2021). We discarded the first minute of data after the respirometers were sealed to allow proper mixing of the water in the chambers. Fish’s SMR estimates were analysed using the FishResp package in R (Morozov et al., 2019). We included all overnight measurements taken from the point where *Ṁ*O2 stabilised (∼6/7 hours). SMR was calculated as the mean of the lowest 20th percentile with an R^2^ for each slope set at 0.95 (Chabot et al., 2016). We then subtracted SMR from MMR to calculate the aerobic scope (AS). We measured background respiration on empty chambers for three 10-minute cycles at the trial’s start and end. We subtracted averaged values from all fish respiration measurements, assuming a linear increase in microbial respiration. To limit bacterial proliferation, we cleaned the all apparatus with hydrogen peroxide (H2O2) and warm water every second day and left to dry in sunlight. At the end of each trial, we removed fish from the chamber and euthanized them with an overdose of 10% eugenol (4 mL L^-1^ water). Fish were immediately measured, weighed to the nearest grams, and frozen at -18°C for later organ sampling and parasite screening.

### Tissue sample preparation and parasite screening

Fish heart, brain, spleen, and gills were dissected on ice to prevent cellular degradation (Stemi DV4, Zeiss, Germany). Each organ was weighed (MSE225S, Sartorius Weighing Company, Germany), diluted at 20 times their wet weight with a buffer solution (100 mM potassium phosphate, 20mM EDTA, pH 8.0; modified protocol from (Hunter-Manseau et al., 2019), homogenised with a polytron (PT1200) and immediately stored at -80°C Following organ removal, we screened each fish for parasites by first counting the total number of encysted black spots (metacercaria) present on the fish body, fins, muscles and gills on both sides of the fish. Prevalence of black-spot infection was 100% with varying intensity (range 9 - 1314 metacercaria cysts) (Fig. 1A). Fish digestive tract, liver and gonads were dissected to screen for other endoparasites. Infection by bass tapeworms (*Proteocephalus ambloplitis*) was primarily found in the liver and digestive tract (prevalence: 99%; range: 0 - 301) (Fig. 1B). 40 fish were also infected by yellow grub trematodes (*Clinostomum marginatum*) found in muscles, heart, gills and brain (prevalence: 33%; range: 0 - 14) (Fig. 1C). We also found nematode larvae in the body cavity of 6 fish (prevalence: 4.95%; range: 0 - 1) (Fig. 1D). These parasites are hereafter grouped under the term internal parasites. Finally, we corrected fish mass by parasites’ mass (Lagrue & Poulin, 2015). We used the mass calculation for bass tapeworm and yellow grub determined by Guitard et al. (2022) and weighed each nematode individually. The fish mass correction did not account for black spots due to their negligible weight (< 0.000001g). For 1 fish acclimated at 25°C and 1 fish acclimated at 30°C, we were not able to precisely quantify the internal parasite intensity due to the extremely high parasite aggregation in their livers. For both fish, we attributed the same number of cestodes as the highest number counted in our sample (301 cestodes), which is an underestimate of the parasite intensity in these individuals.

**Figure 1.**
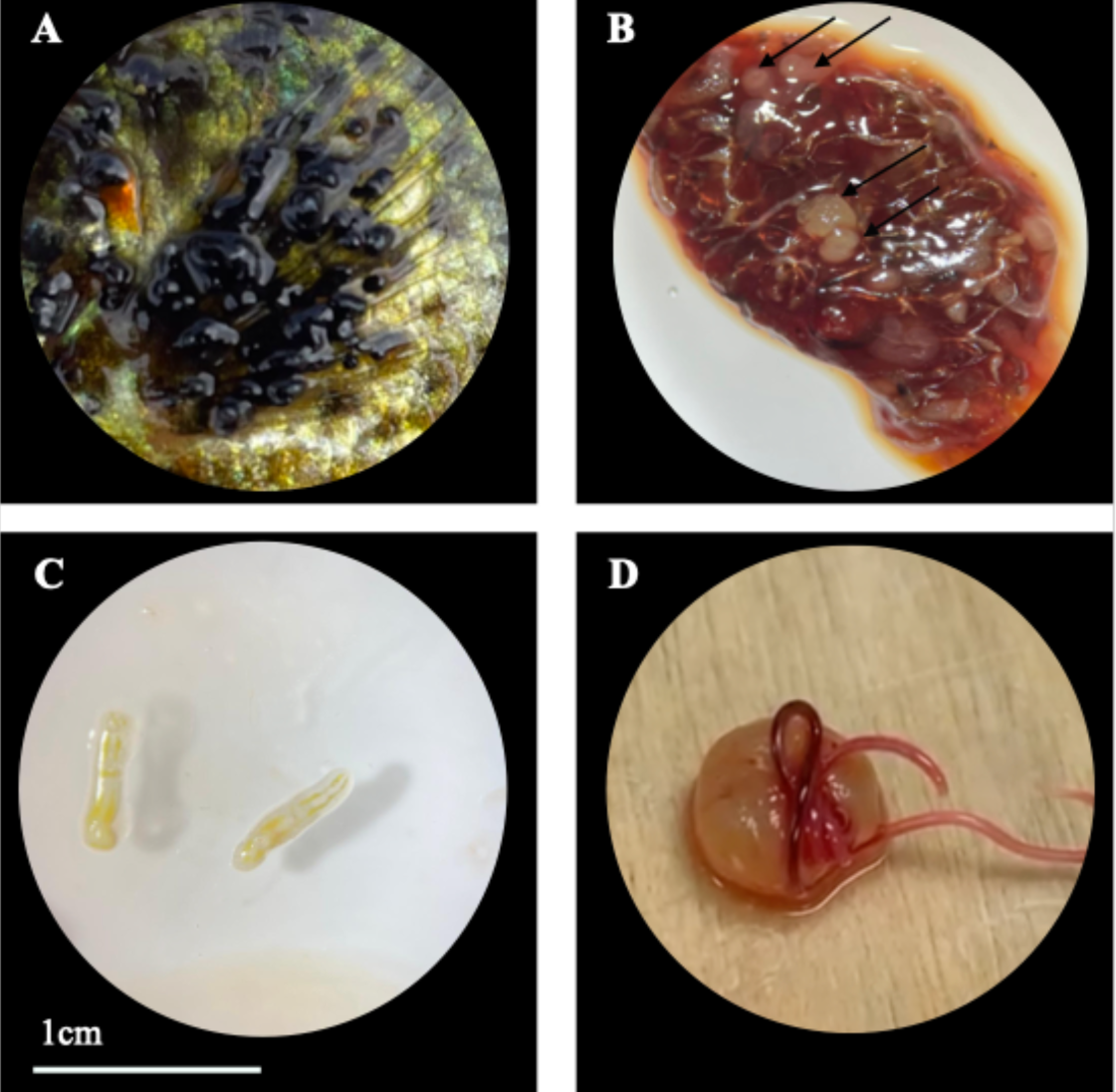
Pictures of the four parasite species found in sampled pumpkinseeds: (A) Black spot trematodes (*Uvulifer sp., Apophallus sp.*) shown on the pectoral fin of a pumpkinseed sunfish. Each black dot represents an individual parasite. (B) Bass tapeworms (*Proteocephalus ambloplitis*) encysted in the liver and pointed at by black arrows. (C) Yellow grub trematodes (*Clinostomum marginatum*). (D) Nematode larvae emerging from its eggshell. Scale (bottom left) applies to all panels of the figure.

### Mitochondrial enzymatic activity

We measured enzymatic activities on 4 key organs (heart, brain, spleen and gills). The cardiovascular system was selected because water temperature is known to strongly affect its mitochondrial functions (Chung, Bryant, and Schulte 2017) while little is known about how parasites may impact these relationships (but see Mélançon et al., 2023). We selected the spleen (lymphoid organ) for its critical role in immune activation in fish infected by helminth parasites (Zapata et al., 2006). Finally, we selected the brain because parasites and temperature can both influence cellular activity (Chung, Bryant, and Schulte 2017; Nadler et al., 2021).

Regarding the enzymes studied, the activity of citrate synthase (CS) was assessed as it is an indicator of citric acid cycle activity (aerobic metabolism). We measured lactate dehydrogenase (LDH) activity, as LDH is used by fish to produce energy via glycolysis (anaerobic pathways) (Soengas & Aldegunde, 2002). Finally, we assayed both the electron transport system complexes I + III and complex IV (cytochrome *c* oxidase; CCO) for their role in oxidative phosphorylation (OXPHOS), a merging point where metabolic pathways interact to support energy production (Blier et al., 2014). We measured enzymes activity via spectrophotometrically using a microplate reader (Mithras LB940 microplate reader, Berthold Technologies, Germany) and its associated software (MikroWin 2010, Labsis Laborsysteme, Germany). Protocols were adapted from Thibault et al. (1997), Hunter-Manseau et al. (2019) and Mélançon et al. (2023). Each measurement was performed in duplicate and in a temperature-controlled room set at the acclimation temperature of the fish. Enzymatic activities were normalised for protein concentration (mg·ml ^-1^), determined with the bicinchoninic acid assay (BCA) method (Smith et al., 1985). Due to the limited amount of tissues for some fish and organs, we were not able to measure all enzymatic activities in all organs. A detailed description of each assay protocol is provided as supplementary material (supportive text and Table S2) and details on replication are shown in Table 1.

**Table 1.** Replication statement for the experimental design examining how parasite and acclimation temperature affects fish metabolism at different biological scales.

|  | **Scale of inference** | **Scale at which the factor of interest is applied** | **Number of replicates at the appropriate scale** |
| --- | --- | --- | --- |
| Whole-organism metabolism | Individuals | Individuals assayed at their acclimation temperature (20°C, 25°C or 30°C) | The level of replication was 40 individuals per acclimation temperature. |
| Cellular-metabolism | Individuals | Organs (heart, brain, spleen and gills) assayed at the acclimation temperature fish were acclimated at. | The level of replication varied per organs and temperature due to limited amount of tissues (for detailed information Table S2). |

### Data analysis

All data were analysed using R v.4.3.1 (R Core Team 2023). First, we used linear models to test whether acclimation temperature and parasite intensity were related to a fish’s whole-organism metabolic traits (MMR, SMR, and AS). All three models had log-transformed metabolic variables as response variables while acclimation temperature, internal parasites, black spot, and log-transformed mass (parasite-corrected fish mass) were fixed effects. We included an interaction between each parasite group and acclimation temperature. Candidate models were selected based on Akaike’s Information Criterion (AIC) (Burnham et al., 2011). In all cases, we kept in the final model the fixed effects acclimation temperature, black spot intensity, internal parasite intensity and fish mass and the interaction between acclimation temperature and internal parasite. We tested variables present in the final models for both collinearity and variance inflation (VIF; car package). We used post hoc tests for pairwise comparison between means of acclimation temperature groups (*emmeans* package; (Lenth, 2018)). A visual inspection of the diagnostic plots ensured that all assumptions were met for each selected model. Second, we used a multivariate linear regression model as our measurements of enzyme activity are highly correlated due to the fact that we took multiple measurements within each organ. We built one model per enzyme measured where we had a matrix of organs (heart, brain, spleen, gills) bound as a multivariate response variable. For each organ, we log transformed enzyme activity. Acclimation temperature, internal parasite, black spot, and the interaction between each parasite group and acclimation temperature were fixed effects. We used the mvIC package (Hoffman, 2022) for model selection. Final models included only the fixed effects acclimation temperature, internal parasite and blackspot intensity for each enzyme normalised for protein content.

## RESULTS

### Whole-organism metabolic rate

There was no significant interaction between acclimation temperatures and internal parasite count on MMR, SMR or AS (Table S2).Acclimation temperature positively affected MMR (*F*_2,112_ = 9.904, *p* < 0.001) with MMR increasing between 20°C and 30°C and between 25°C and 30°C (Fig. 1A; Table S3). Acclimation temperatures also had an effect on SMR (*F*_2,112_ = 6.443, *p* = 0.002; Table S3), which significantly increased from to 20°C to 25°C and subsequently decreased from 25°C to 30°C, resulting in no detectable variation in SMR between 20°C and 30°C (Fig. 1A; Table S3). These changes in MMR and SMR translated into significant differences in AS in relation to acclimation temperature (*F*_2,112_ = 7.15, *p* = 0.001). Fish acclimated at 30°C had significantly higher AS compared to both lower temperatures (Fig. 1A; Table S3). Internal parasite was not related to any of the three metabolic traits (MMR: *F*_1,112_ = 0.11, *p* = 0.740; SMR: *F*_1,112_ = 0.814, *p* = 0.368; AS: *F*_1,112_ = 0.012, *p* = 0.910). However, black spot count negatively correlated with MMR (*F*_1,112_ = 6.321, *p* = 0.013) and AS (*F*_1,112_ = 5,505, *p* = 0.020) (Fig. 1B). All three metabolic traits were positively related to fish body mass (Table S2).

**Figure 1.**
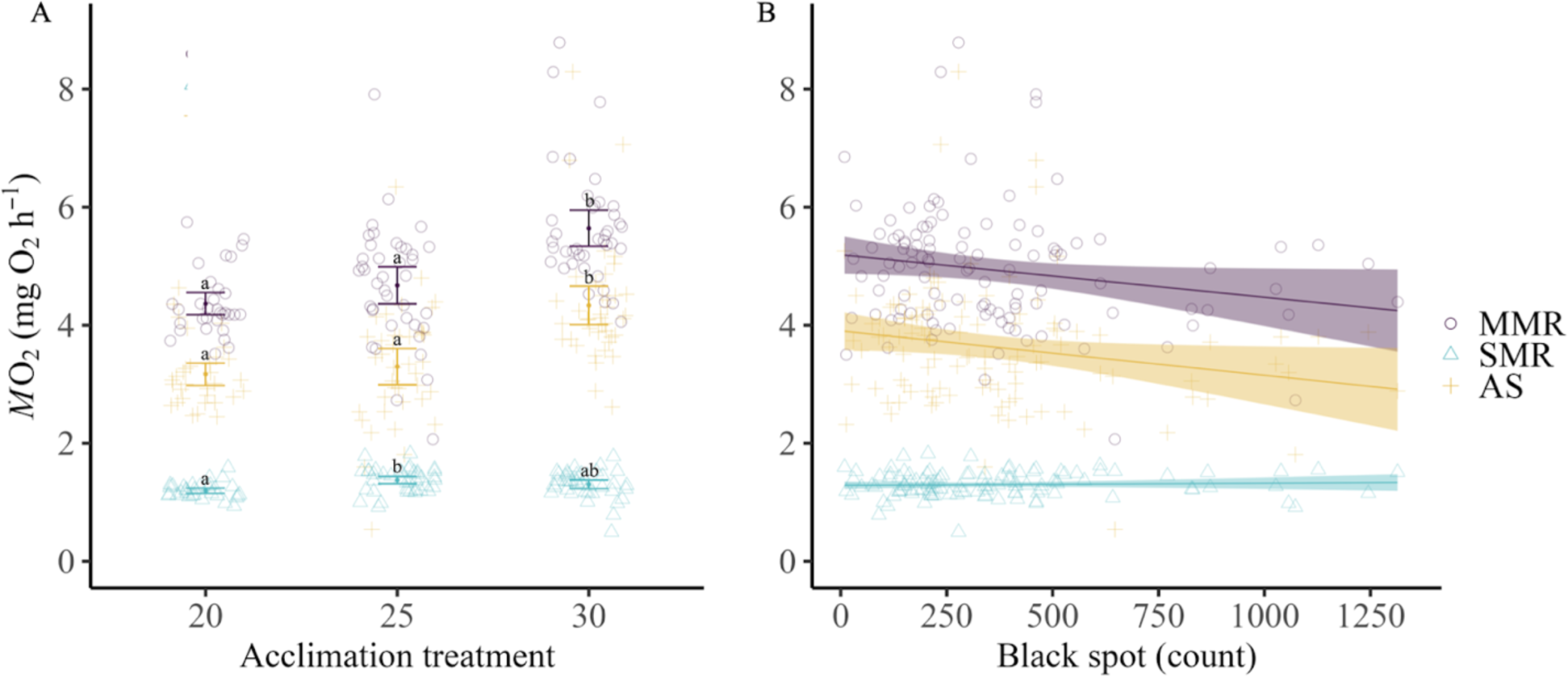
Whole-organism metabolic rates (*Ṁ*O_2_) as a function of (A) acclimation temperature (B) and black spot counts in pumpkinseed sunfish (n = 120). Circles (purple) represent maximum metabolic rate (MMR), triangles (blue) represent standard metabolic rate (SMR), and crosses (yellow) represent aerobic scope (AS). (A) For the relationship between metabolic rates (MMR, SMR, AS) and acclimation temperature, coloured points and error bars represent each acclimation temperature’s mean and confidence interval. Different letters denote significant differences across acclimation temperatures (*p* <0.05). (B) Lines represent linear regressions for each metabolic trait as a function of parasite intensity. The shaded areas represent the upper and lower 95% confidence intervals. All metabolic rates were standardised to the mean fish mass for visual representation (11.46g).

### Mitochondrial enzymatic activity

Acclimation temperature altered maximal activity of enzymes related to aerobic metabolism (CS: *F*_2,80_ = 8.782, *p* <0.001; ETS: *F*_2,89_ =10.780, *p* <0.001; CCO: *F*_2,80_ = 9.343, *p* <0.001) and anaerobic metabolism (LDH: *F*_2,101_ = 18.351, *p* <0.001). For all enzymes, we observed organ-specific patterns in response to acclimation temperature (Fig. 2 & 3).

**Figure 2.**
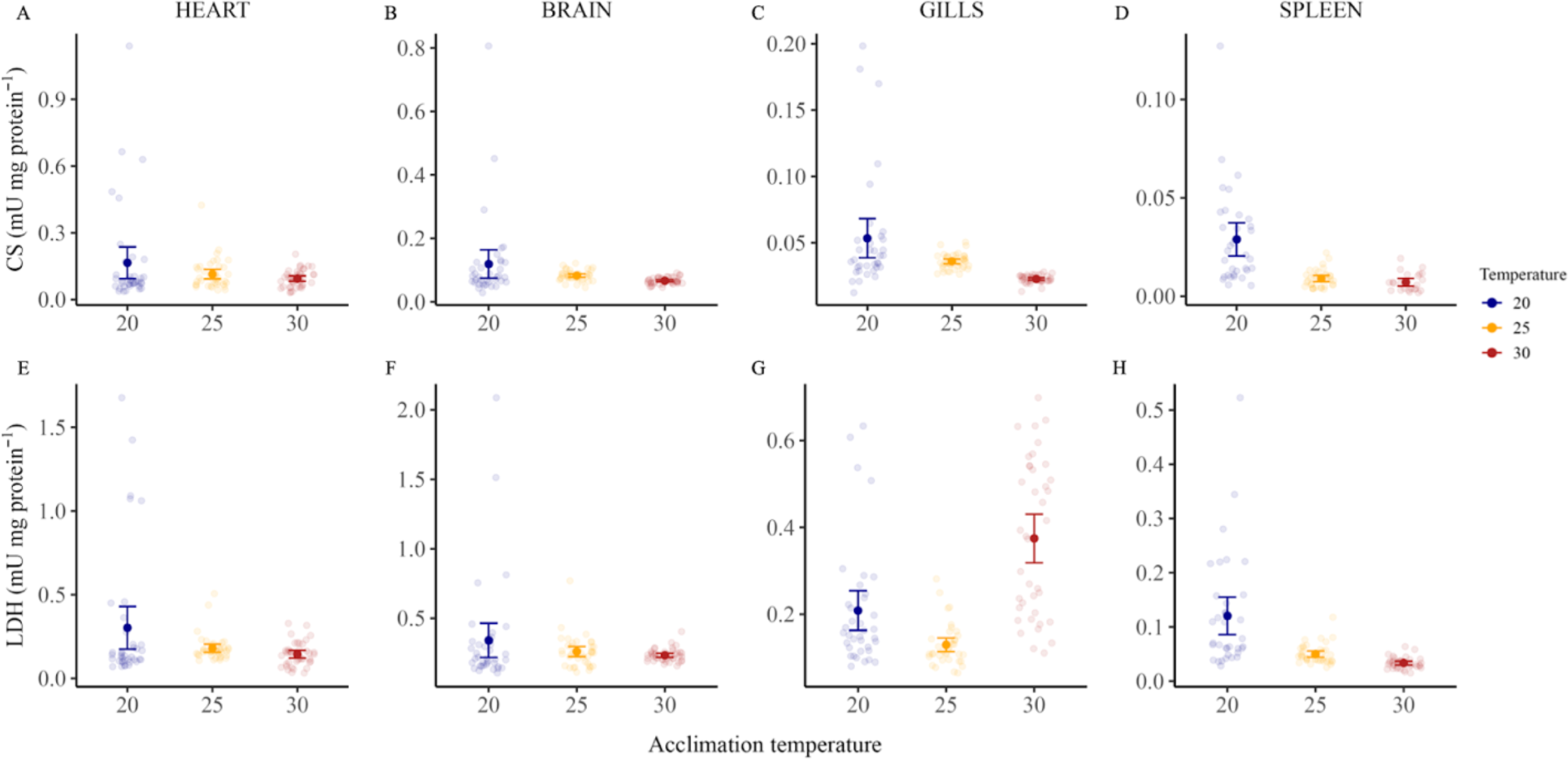
Maximal activity of citrate synthase (CS) and lactate dehydrogenase (LDH) of heart (A, E), brain (B, F), gills (C, G) and spleen (D, H) tissues sampled from pumpkinseed sunfish acclimated at three temperatures. Each panel’s error bars and associated coloured points represent mean and confidence intervals. Each point represents the enzyme activity normalised for protein content of a single fish tested at its acclimation temperature, with blue coloured points representing fish acclimated and tested at 20°C, orange at 25°C and red at 30°C.

All three aerobic enzymes showed no relation between maximal heart activity and acclimation temperature (Fig. 2A-D & Fig. 3; Table S5, S7, S8), while heart LDH anaerobic enzyme activity was altered in response to acclimation temperatures (*F*_2,101_ = 3.093, *p* = 0.049). Despite a negative trend between heart LDH maximal activity and acclimation temperatures, post hoc tests showed a significant difference in means of acclimation temperatures between 20°C and 30°C only (*p* = 0.026) (Fig. 2E).

**Figure 3.**
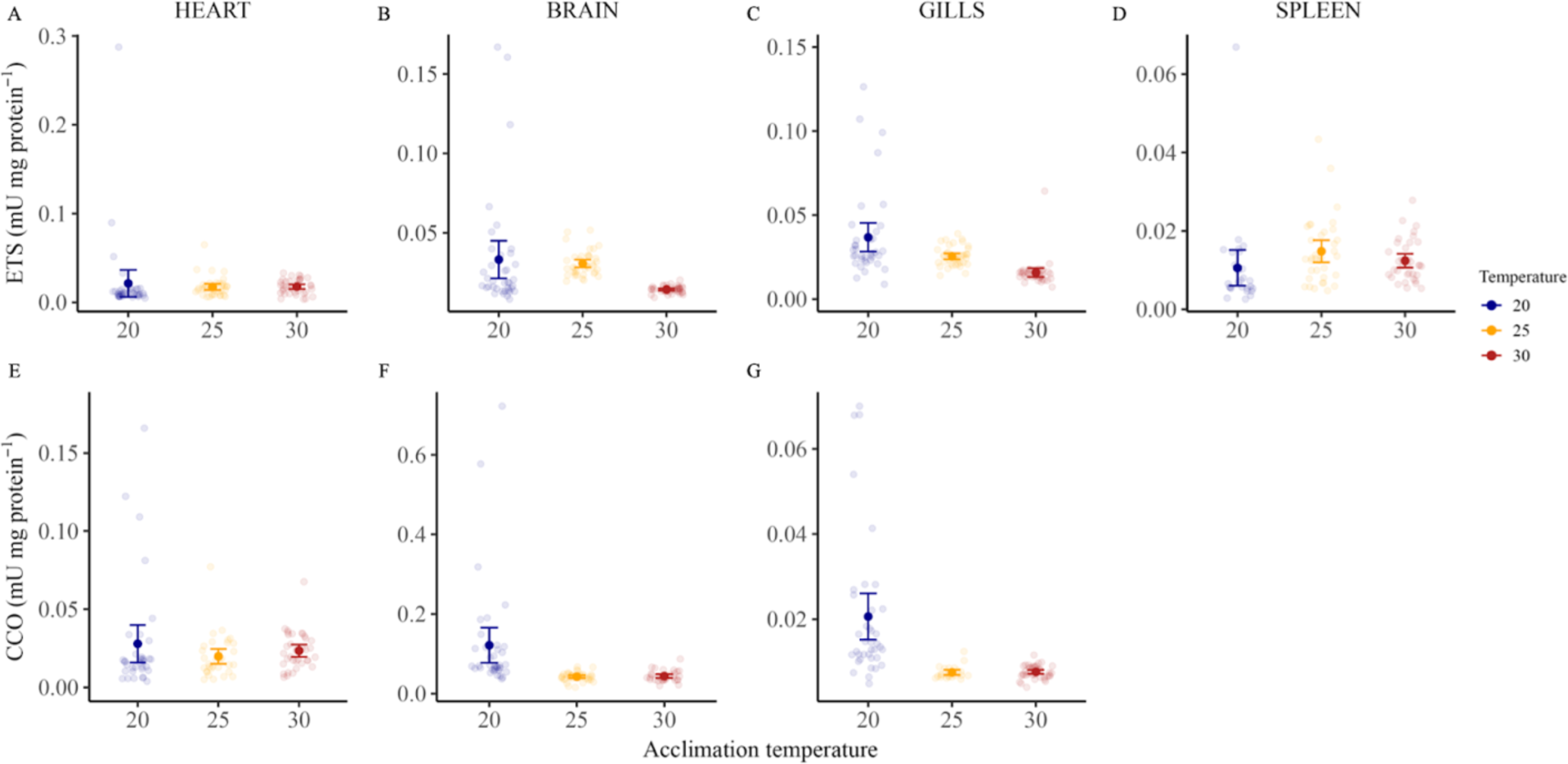
Maximal activity of electron transport system (ETS) and cytochrome *c* oxidase (CCO) of heart (A, E), brain (B, F), gills (C, G) and spleen (D) tissues sampled from pumpkinseed sunfish acclimated at three temperatures. Each panel’s error bars and associated coloured points represent mean and confidence intervals. Each point represents the enzyme activity normalised for protein content of a single fish tested at its acclimation temperature, with blue coloured points representing fish acclimated and tested at 20°C, orange at 25°C and red at 30°C.

Acclimation temperature altered brain CS, CCO and ETS maximal activity (Table S5, S7, S8). Means comparison of acclimation temperatures showed a significant decrease in CS and ETS maximal activity between 20°C and 30°C (CS and ETS: *p* < 0.001) and between 25°C and 30°C (CS: *p* = 0.044; ETS: *p* < 0.001), but not between 20°C and 25°C (CS: *p* = 0.500; ETS: *p* = 0.077) (Fig. 2B and 3B). CCO maximal activity decreased significantly between 20°C and 25°C (*p* < 0.001) and between 20°C and 30°C (*p* < 0.001) but not between 25°C and 30°C (*p* = 0.970; Fig. 3F). Anaerobic enzyme LDH showed no altered activity from acclimation temperature in the brain (Fig. 2F; Table S6).

Spleen ETS maximal activity was significantly altered by acclimation temperature with a significantly decreasing activity at 20°C compared to both 25°C and 30°C (Fig. 3D; Table S7). Both CS and LDH maximal activity showed a significant negative relationship between maximal activity and increased acclimation temperatures (Fig. 2D, H; Table S5 - S6). LDH activity was significantly different among all acclimation treatments (all *p* < 0.001; Fig. 2H), whereas for CS, means comparison showed differences between groups 20°C and 25°C (*p* < 0.001) and between 20°C and 30°C (p < 0.001), but not between 25°C and 30°C *p* = 0.128; Fig. 2D).

Acclimation temperature significantly affected gills CS, ETS and CCO maximal activity (all enzymes: *p* < 0.001; Table S5, S7, S8). Mean comparisons showed lower ETS and CCO enzyme activities between 20°C and 25°C (*p* < 0.05) and between 20°C and 30°C (*p* < 0.001) but not between 25°C and 30°C (ETS: *p* = 0.646; CCO: *p =* 0.994; Fig. 3C and 3F). Gill CS activity decreased with increasing acclimation temperature with significance between 20°C and 30°C (*p* < 0.001), marginal non-significance between 20°C and 25°C (*p* = 0.057), and no significance between 25°C and 30°C (*p* = 0.994; Fig. 2C). Gill LDH maximal activity was also altered by acclimation temperature (*p* < 0.001) with reduced activity between 20°C and 25°C (*p* = 0.003) followed by a significant increase between 25°C and 30°C (*p* < 0.001). This increased activity at 30°C was well above the maximal activity of the 20°C groups, resulting in a significant difference between means of acclimation groups 20°C and 30°C (*p* < 0.001; Fig. 2G).

Black spot intensity and internal parasite intensity was not related to maximal enzyme activity and that across all measured organs (Table S5-S8). However, there was a positive trend between gill LDH maximal activity and internal parasite intensity (*F*_1,101_ =3.665*; p* = 0.058; Table S6).

## DISCUSSION

Our objectives were to determine the effects of acclimation temperature, parasite infection and their interaction on host fish metabolism at the cellular and whole-organism levels. While we found no evidence of interacting effects between acclimation temperature and parasite intensity, thermally acclimated fish showed whole-organism thermal compensation in maintenance metabolism with no change in SMR across acclimation temperatures, but increasing MMR and, AS. Black spot infection showed the opposite pattern, with MMR and AS being negatively correlated with increasing black spot intensity, while there was no detectable link between black spot intensity and SMR. Moreover, we found no evidence of altered cellular metabolism (enzymatic activities) in response to parasite intensity. Acclimation temperature had a significant effect on enzyme activity with a higher activity of enzymes linked to anaerobic metabolism (LDH) compared to ones linked with aerobic energy production (CS, ETS, CCO). Overall, these results emphasise that parasites and temperature, both essential elements of the biotic and abiotic environment, can have differing effects on components of the metabolic phenotypes of fish at different biological scales.

### Physiological consequences of parasite infection

We found no evidence for a relationship between SMR and blackspot infection (Fig. 1B). Although a study have found trends between SMR and black spot intensity (Thambithurai et al., 2022), our results align with other studies suggesting that established trematode infection has no measurable metabolic costs to hosts (du Toit et al., 2024; Guitard et al., 2022; Nadler et al., 2021). Contrary to SMR, there was a negative relationship between black spot infection and MMR as well as AS (Fig. 1B). Fish in our study were naturally infected and thus we cannot establish a causal link between parasite intensity and the metabolic traits measured. Fish with low MMR and AS may be more susceptible to infection since these traits are associated with reduced swimming capacity (Norin & Clark, 2016). Indeed, wood frogs *Lithobates sylvaticus* spending less time swimming were more susceptible to infection from ranavirus (family: Iridoviridae) (Araujo et al., 2016). On the other hand, infection can induce lethargy in hosts as immune activation increases energy demands (Levet et al., 2024; Lopes et al., 2021). Thus, infection itself may cause decreased metabolic performance as a result of altered energetic allocation. A recent study on pumpkinseed suggests that individual movement behaviours are both predictors of infection susceptibility and are altered following infection suggesting the complicated nature of host-parasite effects on behaviour and physiology (Gradito et al., *In press*). Further work quantifying metabolic traits pre- and post-natural infection is needed to establish causality and determine adaptive changes of metabolic phenotype in response to infection.

Internal parasite intensity was not related to any of the metabolic traits measured at the whole-organism level (Table S2). The level of infestation in our study (mean: 54 parasites per fish) exceeds the intensity observed previously in similar-sized pumpkinseed from the same population (mean: 12 parasites per fish), where metabolic rates were found to decrease in response to parasite intensity (Guitard et al., 2022). This discrepancy may be explained by the amount of time fish spent in the laboratory environment (3-5 days vs. 1 month in this study). Indeed, physiological differences between wild and captive animals are well-known (Turko et al., 2023). Any physiological effects induced by infection may have been masked in our fish by the time they were tested. Therefore, it is possible that our results diverged from previously cited studies because the host’s energy demands varied in response to holding conditions. Further investigations should be made to determine how host energy demands change over time in captive parasitized fish.

Similar to results described in Mélançon et al. (2023), we observed no differences in maximal enzyme activity in relation to either black spots or internal parasite intensity (Tables S3 - S6). However, some parasites have been shown to impact cellular metabolism: brain-infecting trematode *Euhaplorchis californiensis* alter California killifish *Fundulus parvipinnis* neuronal signalling through an effect on LDH levels in the brain (Nadler et al., 2021). Futhermore, recent work on several populations of pumpkinseed found that cestode density increased maximal LDH activity in the liver at the location of the infection (Sabbagh et al., *In press*). We included neither the liver nor the muscles (both sites of parasite infection) in our study to avoid the risk of measuring enzyme activity coming from the parasites. However, a recently developed protocol testing enzyme activity before and after the removal of parasites from infected organs showed no evidence for contamination in fish enzyme activity from parasite tissue (Pepin et al., unpublished data). Therefore, a future direction could be to integrate measurements of organs where parasite infection is located as the cellular response may differ in those organs because of inflammation, tissue dysfunction and necrosis caused by parasite presence.

### Aerobic performance across acclimation temperatures

Aerobic scope (AS) significantly increased across acclimation temperature as a result of increased MMR and no detectable changes in minimum energy demands (SMR) (Fig. 1A). Such results indicate that fish were able to compensate across thermal regimes. A recent study demonstrated that the critical thermal limit of pumpkinseed from the same population as the one studied here averaged 35.58°C, 38.66°C and 40.65°C after three weeks of acclimation at 20°C, 25°C, and 30°C respectively (De Bonville, Côté, and Binning 2024). Thus, the metabolic traits were measured on our fish at temperatures well below their critical thermal limit. This may explain why at temperatures above their natural thermal range (30°C), they were able to counteract the direct effect of elevated temperature on metabolism and sustain aerobic performance (Fig. 1A). Chronic exposure to elevated temperature can cause a state of physiological stress which ultimately compromises fish fitness and survival (Alfonso et al., 2021). Thus, it is possible that the thermal compensation observed in our study may come at the cost of decreased growth rate or reproductive success despite fish being well-below their critical thermal limit. Due to the potential implication for fish population dynamics, future work should consider integrating chronic exposure to elevated temperature with co-occuring environmental stressors to better understand physiological changes that impair survival in nature.

Sub-organismal levels of biological organisation are more directly influenced by thermodynamic laws, and our results found evidence for altered maximal activity for enzymes related to aerobic pathways, namely CS, ETS, and CCO (Fig. 2A-D and Fig. 3A-H). Long-term exposure to acclimation temperatures can reduce enzyme activities in the tricarboxylic acid (TCA) cycle, alter glycolysis, and lower enzyme sensitivity to temperature change (Ekström et al., 2017; Pichaud et al., 2019). CS is part of the TCA cycle, which is a central aerobic pathway for producing ATP, while CCO is assumed to be a key regulator of electron flow and thus to control OXPHOS capacity in ectotherms (Blier et al., 2014). As acclimation temperature increased, mitochondrial density decreased in the brain, spleen and gills, reducing the number of mitochondria available to carry out OXPHOS. Downregulation of enzyme activity in response to increasing temperature may indicate organisms’ capacity for metabolic reorganisation by optimising energy production to sustain energetic demands as observed at the whole-organism level (Fig 1A). It can also be a sign of reduced efficiency as temperature can alter substrate availability and, thus, the catalysing rate of enzymes. Furthermore, it may reflect a shift in metabolic pathways away from aerobic metabolism and towards alternative energy-producing pathways suggesting that acclimation causes a shift in the way energy is produced.

Fish held at 30°C had significantly higher gill LDH activity compared to lower acclimation temperatures, suggesting a strong reliance on anaerobic metabolism to produce energy at elevated temperatures. Warm acclimation can lead to changes in gills morphology (Johansen et al., 2021). For instance, prolonged exposure to elevated temperature can alter oxygen diffusion across the lamellae, the amount of interlamellar mass, which ultimately modifies the total surface of respiratory lamellae gills (Bowden et al., 2014). Such adaptation can lead to prolonged increases in LDH activity. For example, several weeks are needed for yellowtail fusilier (*Caesio cuning)* to return to LDH activity levels similar to pre-acclimation ones (Johansen et al., 2021). Organs have a time-dependent response to acclimation and may adjust to acclimation temperature at a different pace (Bouchard & Guderley, 2003). Future work on thermal acclimation should consider this time-course aspect if we want to fully comprehend how organisms adapt and adjust to thermal variation in nature.

## CONCLUSION

Our findings suggest that temperature and parasite infection impact the metabolism of pumpkinseed sunfish, with varying effects at different biological levels. Acclimation temperatures affected metabolic rate estimates, with aerobic scope maintained due to increased maximum metabolic rate (MMR) without significant changes in standard metabolic rate (SMR). This suggests fish can sustain aerobic performance even at temperatures above their natural range, which may be advantageous in the context of global climate change. Altered maximal activity of all enzymes in response to acclimation temperature indicated that metabolic changes at sub-organismal levels may have contributed to thermal compensation at the whole-organism level. In contrast, parasite infection was not related to enzyme activity, while increased black spot intensities were related to reduced MMR and AS, suggesting compromised aerobic capacity in parasitized fish. Despite no interaction between stressors, our findings illustrate that temperature and parasite infection can each disrupt metabolism, affecting fish performance. This highlights the ecological importance of understanding how multiple environmental stressors independently and collectively affect freshwater organisms.

