## Supplementary material for "Acclimation temperature and parasite infection drive metabolic changes in a freshwater fish at different biological scales"

*Enzyme activity preparation and measurements*

**Citrate synthase** (CS) activity was estimated by the rate of TNB formation (ε412 = 14.15 $mM^{-1} \cdot cm^{-1}$) measured at 412nm over 7 minutes. The reaction medium was 100 mM imidazole-HCl, 0.1 mM DTNB, 0.1 mM acetyl-CoA, 0.15 mM oxaloacetic acid, pH 8.0. We measured the specificity of the reaction in the absence of oxaloacetic acid for all samples, followed by the final assay.

**Lactate dehydrogenase** (LDH) activity was estimated as the rate of NADH oxidation (ε340 = 6.22 $mM^{-1} \cdot cm^{-1}$), measured at 340 nm for 5 minutes. The reaction medium was 100mM potassium phosphate, 0.16mM NADH, 0.4 mM pyruvate (omitted for the control), pH 7.0.

**Electron transport system (ETS - mitochondrial complex I and II**) activity was estimated as the rate of p-iodonitrotetrazolium violet (INT) reduction (ε490 = 15.9 $mM^{-1} \cdot cm^{-1}$), measured at 490 nm for 5 minutes. The reaction medium was 100 mM potassium phosphate, 0.85 mM NADH, 2 mM INT and 0.03% (v/v) triton X 100, pH 8.5. We verified the specificity of the reaction with two controls: one without sample and one in the absence of NADH (CI-substrate)

**Cytochrome *c* oxidase** (CCO) activity was estimated as the rate of cytochrome c oxidation (ε550 = 18.5 $mM^{-1} \cdot cm^{-1}$), measured at 550 nm for 5 minutes. The reaction medium was 100 mM potassium phosphate, 0.05% (v/v) tween-20, 50 µM cytochrome c, 1 mM ADP, 0.03% (v/v) triton

X 100- and 4.5-mM dithionite, pH 8.0. The solution was bubbled for 5 minutes, followed by an absorbance ratio (550:565 nm) measurement with a threshold set at 9. If the absorbance ratio happened to be below 9, we discarded the solution. We verified the specificity of the reaction with 40 mM sodium azide (CCO-inhibitor).

### TABLES AND FIGURES

Table S1 — Detailed information of the intermittent flow-respirometry set-up used to estimate fish's standard metabolic rate (SMR), maximum metabolic rate (MMR) and background respiration. The following checklist is based on guidelines from Killen et al. (2021).

| Detailed information | Fish status during measurements |
|---|---|
| Fish body mass | 11.67 ± 2.93g (mean ± SD) |
| Volume of empty respirometer including recirculation tubes | 490 mL |
| Equipment to achieve mixing in the chamber | We used a recirculating pump (Decdeal 2.4W, 250l $h^{-1}$), and each chamber had baffles on each end to diffuse the water being pumped around. |
| Material of tubing used in the mixing circuit | Tygon |
| Declare whether volume of tubing in mixing circuit was included in calculations of oxygen uptake | Yes, included |
| Material of the respirometry chambers | Acrylic |
| Type of oxygen probe and data recording | FireSting$O_2$ Optical Oxygen meter from Pyro Science GmbH, Aachen, Germany |

| | |
|---|---|
| Method of probe calibration | Two-point calibration (100% air saturation and 0% air saturation with sodium sulfite) at the beginning of each trial. |
| Sampling frequency of oxygen concentration in water | 3s |
| Oxygen probe placement | Placed in the recirculation circuit |
| Flow rate during flushing and recirculation, or confirm that chamber returned to normoxia during flushing | Yes, each chamber returned to normoxia during the flushing period |
| Timing of each measurement cycle (i.e., flush/closed cycles) | 4 minutes of flush / 6 minutes of closed cycles with cycle control by a digital timer (T319, Permanent Industry Co. China) connected to respirometer chamber pump. |
| State whether software temperature compensation was used during recording of water oxygen concentration | Yes, we used it |
| Temperature during the respirometry trial | 20°C ; 25°C and 30°C |
| How temperature was controlled | Hakke A10, SC100 (Thermo Scientific™) water chiller / heater system. |
| Photoperiod during respirometry | 14h:10h (day/night cycle) |
| Describe if ambient water bath was cleaned and aerated during measurement of oxygen uptake, and if so, how this was done (e.g. filtration, periodic or continuous water changes) | We used a closed system to ensure stable water temperature. Water was UV sterilized throughout the trial. We had a continuous water exchange between the water bath holding the chambers and the water tank. We used an Eheim Compact 300 pump (EHEIM; GmbH & Co., Germany) to pump water in and out of each bath. |
| Describe whether chambers were visually shielded from external disturbance | Chambers were covered in opaque PVC material with a small viewing window on the top. |
| The number of fish measured during a given respirometry trial | 4-12 |

| | | |
|---|---|---|
| Fish were able to see each other during measurements? | No | |
| Duration of animal fasting before placement in respirometer | 24h | |
| Duration of all trials combined (number of days to measure all animals in the study) | 15 days. We tested the first batch of fish on July 4th. 2022, and the last on July 19th 2022. | |
| Acclimation time to the laboratory (or since capture) before respirometry measurements | Minimum 28 days since capture | |
| Was background respiration measured? | Yes | |
| Method used to account for background respiration | We measured it on empty chambers at the beginning and the end of each trial. We used three cycles of 10 minutes. | |
| How were changes in background respiration modelled over time? | linear | linear |
| Level of background respiration (e.g., as a percentage of SMR) | < 0.1% | < 0.1% |
| Method and frequency of system cleaning | The system was cleaned with a mix of warm water and hydrogen peroxide ($H_2O_2$) every second day. The water was sterilised with UV lamps during the trial (i.e., metabolic measurements) | |
| Time excluded from closed measurement cycles for estimation of SMR/ lowest metabolic rate | Time needed for to $\dot{M}O_2$ be stable (on avergae 5 hours after being placed in the chamber) | |
| Duration over which metabolic rate was estimated | ~ 8 hours | |
| What value was taken as SMR (pre-injection) and lowest metabolic rate (post-injection)? | We took the lowest 20th percentile for estimating SMR (Chabot, Steffensen and Farrell, 2016) | |
| When was MMR measured in relation to SMR? | Before SMR measurement | |
| Method used to measure MMR | Manual chase to exhaustion for 3 minutes | |

| | |
|---|---|
| Was air exposure added after exercise? | Yes, we added 1 minute of air exposure |
| Value taken as MMR's | Highest rate of oxygen uptake over a two-minute rolling interval (Prinzing et al., 2021) |
| Duration of the slopes used to calculate MMR | ~ 15 min |
| Slope estimation used to calculate MMR | Rolling regression |
| Calculation of absolute aerobic scope | Substraction of SMR absolute value from MMR ones. |

Table S2 — Detailed information summarising the enzymatic assays performed for each sample, organ and acclimation temperature.

| | | **20°C** | **25°C** | **30°C** |
|---|---|---|---|---|
| CS | | | | |
| | Heart | *n* = 38 | *n* = 39 | *n* = 40 |
| | Brain | *n* = 37 | *n* = 39 | *n* = 40 |
| | Spleen | *n* = 33 | *n* = 34 | *n* = 25 |
| | Gills | *n* = 35 | *n* = 40 | *n* = 37 |
| LDH | | | | |
| | Heart | *n* = 38 | *n* = 40 | *n* = 40 |

| | | | |
|---|---|---|---|
| Brain | *n* = 38 | *n* = 38 | *n* = 40 |
| Spleen | *n* = 35 | *n* = 40 | *n* = 38 |
| Gills | *n* = 37 | *n* = 37 | *n* = 39 |
| ETS | | | |
| Heart | *n* = 37 | *n* = 38 | *n* = 38 |
| Brain | *n* = 38 | *n* = 39 | *n* = 38 |
| Spleen | *n* = 27 | *n* = 36 | *n* = 37 |
| Gills | *n* = 37 | *n* = 40 | *n* = 39 |
| CCO | | | |
| Heart | *n* = 35 | *n* = 32 | *n* = 36 |
| Brain | *n* = 38 | *n* = 38 | *n* = 32 |
| Gills | *n* = 38 | *n* = 23 | *n* = 40 |

Table S3 — Whole-organism metabolic rates ($\dot{M}O_2$) at the three acclimation temperatures, adjusted to an overall mean body mass of 11.46 g

| | **20°C** | **25°C** | **30°C** |
|---|---|---|---|

| | | | |
|---|---|---|---|
| *n* | 40 | 40 | 40 |
| MMR (mg $O_2 h^{-1}$ ) | 4.38 ± 0.08$^a$ | 4.68 ± 0.16$^a$ | 5.64 ± 0.15$^b$ |
| SMR (mg $O_2 h^{-1}$ ) | 1.19 ± 0.0227$^a$ | 1.38 ± 0.03$^a$ | 1.31 ± 0.03$^b$ |
| AS (mg $O_2 h^{-1}$ ) | 3.19 ± 0.08$^a$ | 3.30 ± 0.16 $^b$ | 4.34 ± 0.16$^{ab}$ |

Sample size for each acclimation temperature is denoted by *n*. MMR: maximum metabolic rate, SMR: standard metabolic rate, AS: aerobic scope for whole-organism metabolic rates. All values are given as mean ± s.e.m. Subscript lower case letters indicates significantly different groups.

Table S4 — Results of linear models testing the effect of acclimation temperature (20°C, 25°C, 30°C) and parasite (black spot /internal parasite intensity) and their interaction on SMR, MMR, and AS in pumpkinseed sunfish. Each model was selected by Akaike's Information Criterion (AIC).

| **Response** | **Predictors** | **d.f.** | ***F*-values** | ***P*-value** | ***$R^2$*** |
|---|---|---|---|---|---|
| MMR (mg $O_2 h^{-1}$ ) | Body mass | 1,112 | 125.541 | **< 0.001** | 0.54 |
| | Acclimation temperature | 2,112 | 9.904 | **< 0.001** | |
| | Black spot | 1,112 | 6.321 | **0.013** | |

| | | | | | |
|---|---|---|---|---|---|
| | Internal parasite | 1,112 | 0.110 | 0.740 | |
| | Internal parasite x acclimation temperature | 2,112 | 0.091 | 0.913 | |
| SMR (mg $O_2 h^{-1}$) | Body mass | 1,112 | 128.225 | **< 0.001** | 0.49 |
| | Acclimation temperature | 2,112 | 6.44 | **0.002** | |
| | Black spot | 1,112 | 0.007 | 0.933 | |
| | Internal parasite | 1,112 | 0.814 | 0.368 | |
| | Internal parasite x acclimation temperature | 2,112 | 0.720 | 0.488 | |
| AS (mg $O_2 h^{-1}$) | Body mass | 1,112 | 51.908 | **< 0.001** | 0.42 |
| | Acclimation temperature | 2,112 | 7.150 | **< 0.001** | |
| | Black spot | 1,112 | 5.505 | **0.020** | |
| | Internal parasite | 1,112 | 0.012 | 0.910 | |
| | Internal parasite x acclimation temperature | 2,112 | 0.031 | 0.968 | |

Value in bold indicates significance $p < 0.05$

Table S5 — Results of multivariate multiple linear regression models analysis testing the effect of acclimation temperature (20°C, 25°C, 30°C) and parasite (blackspot and internal parasite) on citrate synthesis (CS) maximal activity in pumpkinseed sunfish

| **Enzyme activity** (u/mg protein) | **Predictors** | **d.f.** | ***F*-values** | ***P*-value** |
|---|---|---|---|---|
| Heart | Acclimation temperature | 2,80 | 1.020 | 0.365 |
| | Black spot | 1,80 | 1.502 | 0.223 |
| | Internal parasite | 1,80 | 2.162 | 0.145 |
| Brain | Acclimation temperature | 2,80 | 3.712 | **0.028** |
| | Black spot | 1,80 | 0.035 | 0.850 |
| | Internal parasite | 1,80 | 0.055 | 0.815 |
| Spleen | Acclimation temperature | 2,80 | 27.008 | < **0.001** |
| | Black spot | 1,80 | 0.000 | 0.991 |
| | Internal parasite | 1,80 | 2.022 | 0.158 |
| Gills | Acclimation temperature | 2,80 | 19.855 | < **0.001** |
| | Black spot | 1,80 | 0.017 | 0.896 |
| | Internal parasite | 1,80 | 1.176 | 0.281 |

Value in bold indicates significance $p < 0.05$

Table S6 — Results of multivariate multiple linear regression models analysis testing the effect of acclimation temperature (20°C, 25°C, 30°C) and parasite (blackspot and internal parasite) on lactate dehydrogenase (LDH) maximal activity in pumpkinseed sunfish

| **Enzyme activity** (u/mg protein) | **Predictors** | **d.f.** | ***F*-values** | ***P*-value** |
|---|---|---|---|---|
| Heart | Acclimation temperature | 2,101 | 3.056 | 0.051 |
| | Black spot | 1,101 | 0.659 | 0.418 |
| | Internal parasite | 1,101 | 2.114 | 0.149 |
| Brain | Acclimation temperature | 2,101 | 0.392 | 0.676 |
| | Black spot | 1,101 | 0.459 | 0.499 |
| | Internal parasite | 1,101 | 0.084 | 0.771 |
| Spleen | Acclimation temperature | 2,101 | 42.689 | **< 0.001** |
| | Black spot | 1,101 | 1.613 | 0.207 |
| | Internal parasite | 1,101 | 1.110 | 0.294 |
| Gills | Acclimation temperature | 2,101 | 36.192 | **< 0.001** |
| | Black spot | 1,101 | 0.014 | 0.904 |

| | | | | |
|---|---|---|---|---|
| | Internal parasite | 1,101 | 3.665 | 0.058 |

Value in bold indicates significance p < 0.05

Table S7 — Results of multivariate multiple linear regression models analysis testing the effect of acclimation temperature (20°C, 25°C, 30°C) and parasite (blackspot and internal parasite) on electron transport system complexes I + III ( ETS I + III) maximal activity in pumpkinseed sunfish

| **Enzyme activity** (u/mg protein) | **Predictors** | **d.f.** | ***F*-values** | ***P*-value** |
|---|---|---|---|---|
| Heart | Acclimation temperature | 2,89 | 1.065 | 0.348 |
| | Black spot | 1,89 | 1.176 | 0.280 |
| | Internal parasite | 1,89 | 0.484 | 0.488 |
| Brain | Acclimation temperature | 2,89 | 23.812 | **< 0.001** |
| | Black spot | 1,89 | 0.013 | 0.908 |
| | Internal parasite | 1,89 | 0.751 | 0.388 |
| Spleen | Acclimation temperature | 2,89 | 5.265 | **0.006** |
| | Black spot | 1,89 | 1.192 | 0.277 |
| | Internal parasite | 1,89 | 1.957 | 0.165 |

| | | | | |
|---|---|---|---|---|
| Gills | Acclimation temperature | 2,89 | 34.082 | **< 0.001** |
| | Black spot | 1,89 | 0.050 | 0.822 |

Value in bold indicates significance $p < 0.05$

Table S8 — Results of multivariate multiple linear regression models analysis testing the effect of acclimation temperature (20°C, 25°C, 30°C) and parasite (blackspot and internal parasite) on electron transport system complex IV (cytochrome *c* oxidase; CCO) maximal activity in pumpkinseed sunfish

| **Enzyme activity** (u/mg protein) | **Predictors** | **d.f.** | ***F*-values** | ***P*-value** |
|---|---|---|---|---|
| Heart | Acclimation temperature | 2,80 | 0.955 | 0.389 |
| | Black spot | 1,80 | 0.960 | 0.330 |
| | Internal parasite | 1,80 | 0.077 | 0.781 |
| Brain | Acclimation temperature | 2,80 | 24.772 | **< 0.001** |
| | Black spot | 1,80 | 0.399 | 0.529 |
| | Internal parasite | 1,80 | 0.020 | 0.887 |
| Gills | Acclimation temperature | 2,80 | 30.341 | **< 0.001** |

| | | | |
|---|---|---|---|
| Black spot | 1,80 | 0.488 | 0.486 |
| Internal parasite | 1,80 | 1.319 | 0.254 |

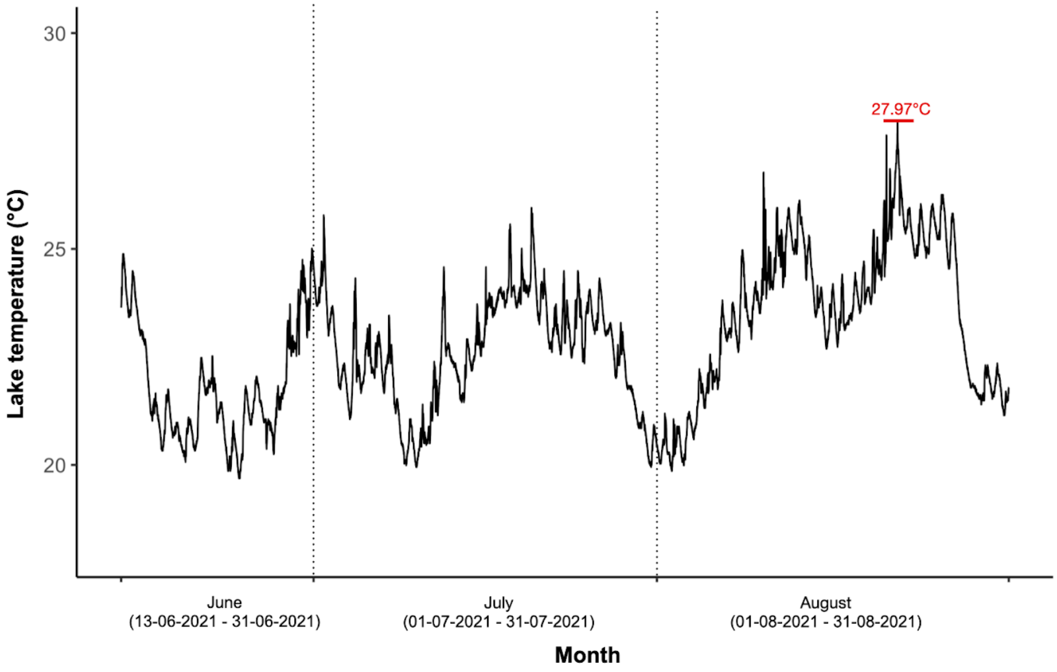


**Figure S1. Log of water temperatures from Lake Cromwell (°C) taken with a HOBO temperature logger (HOBO Pendant MX2201) placed at a 1-meter depth below the lake surface.** Water temperature data were collected from 13-06-2021 to 01-09-2021 with a 10-minute interval between temperature readings. The red line highlights the highest temperature recorded in the lake during the summer of 2021 (27.97°C).

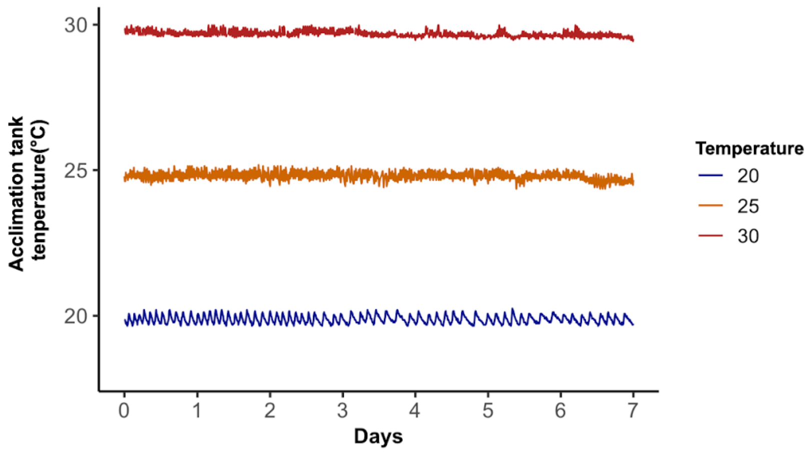


**Figure S2. Log of water temperature from each acclimation tank over one week, recorded with a HOBO temperature logger (HOBO Pendant MX2201) placed inside the tank.** Data displayed were recorded over a seven-day period. Each logger took readings at a 10-minute interval. Each line represents the temperature in one tank: 20°C: in blue, 25°C: in orange, 30°C: in red.
